# Global profiling of the proteome and acetylome in mice with abdominal aortic aneurysms

**DOI:** 10.1101/2025.06.02.657536

**Authors:** Luyao Zhang, Bo Yang, Tao Ding, Yaling Zhao, Yehong Yang, Xiaoyue Tang, Yue Wu, Qiaochu Wang, Zhiyi Zhang, Chunmei Shi, Rong Han, Xutong Zhang, Jiangfeng Liu, Juntao Yang

**Affiliations:** State Key Laboratory of Common Mechanism Research for Major Diseases, Department of Biochemistry and Molecular Biology, Institute of Basic Medical Sciences, Chinese Academy of Medical Sciences & Peking Union Medical College, Beijing, 100005, China; Department of Blood Transfusion, Peking Union Medical College Hospital, Chinese Academy of Medical Sciences, Beijing, 100730, China

**Keywords:** Abdominal Aortic Aneurysm, lysine acetylation, LC-MS/MS, histone, sirtuin protein family

## Abstract

**Background:** Abdominal Aortic Aneurysm (AAA) is a life-threatening vascular condition characterized by a progressive dilation of the abdominal aorta and a high risk of rupture. Currently, no effective pharmacological therapies are available, and treatment relies mainly on surgical intervention. The lack of drug targets is largely due to limited understanding of the molecular mechanisms underlying AAA development. This study aims to identify potential therapeutic targets by performing comprehensive proteomic and acetylomic analyses in a mouse model of AAA.

**Methods:** We conducted global proteomic and acetylomic profiling on abdominal aortic tissues collected from an AAA mouse model. A total of 7,858 proteins and 1,790 acetylated proteins encompassing 4,581 acetylation sites were quantified. Bioinformatics analyses were performed to investigate the regulatory networks and biological pathways associated with these proteins and post-translational modifications.

**Results:** Integrated analysis revealed that histone-related proteins were significantly enriched in pathways co-regulated by the proteome and acetylome. These pathways are influenced by histone acetyltransferases and deacetylases, implicating histone acetylation in AAA-related inflammatory and immune processes. Notably, the deacetylases Sirt2 and Sirt5 were identified as potential regulators of neutrophil extracellular trap (NET) formation through suppression of histone acetylation.

**Conclusions:** This study provides the first comprehensive proteomic and acetylomic dataset for abdominal aortic tissues in a mouse model of AAA, substantially expanding the mouse acetylation database. The results highlight the central role of histone modifications in immune dysregulation and inflammation, and propose Sirt2 and Sirt5 as candidate therapeutic targets that may inhibit NET formation and slow AAA progression. These findings suggest new avenues for drug development targeting the sirtuin family, although further experimental validation and cross-species studies are needed to confirm translational relevance.

**Graphic Abstract:** 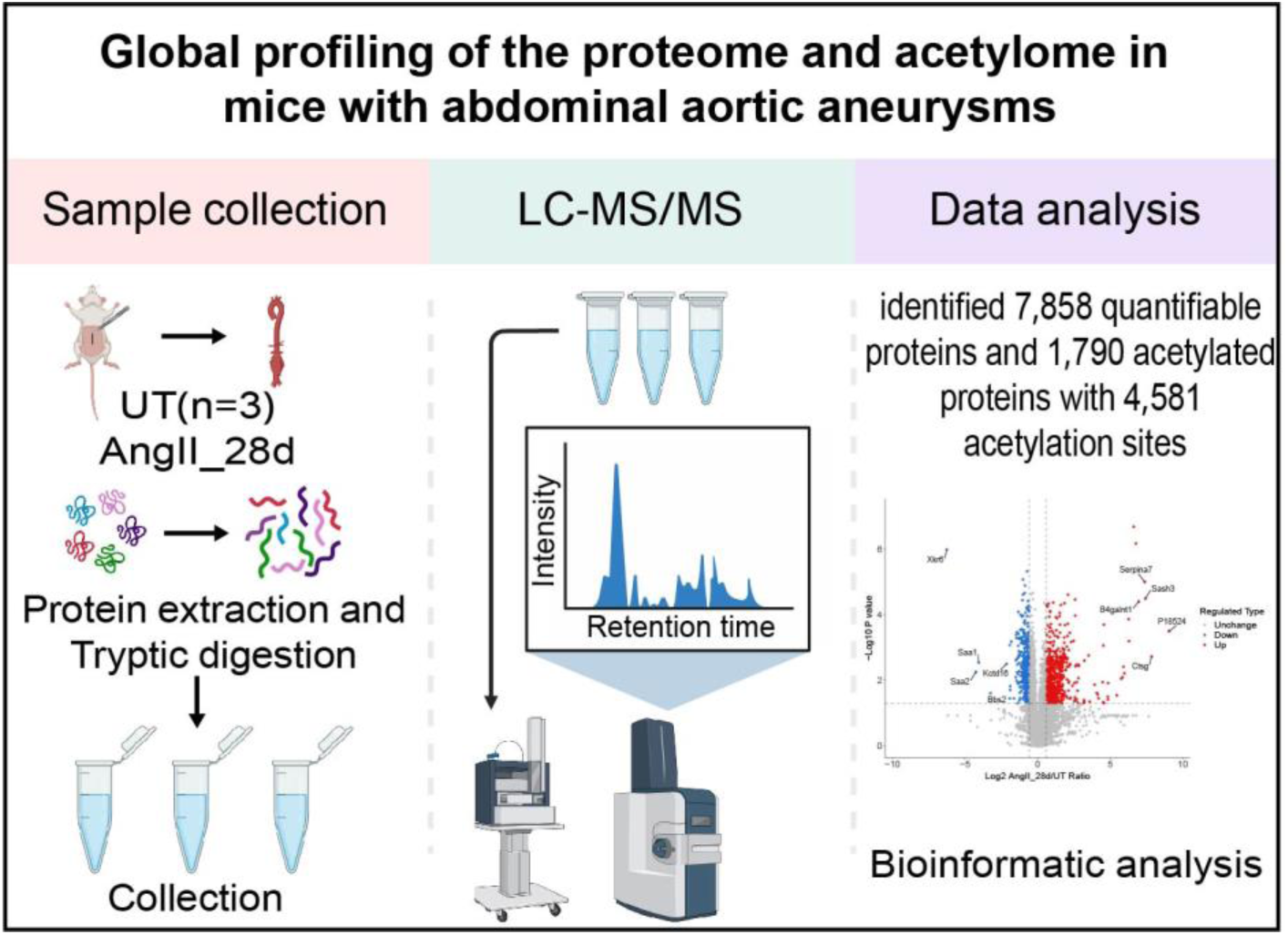

## 1 Introduction

Abdominal Aortic Aneurysm (AAA) ^1^is a common vascular disease characterized by localized dilation and weakening of the aortic wall, which can ultimately lead to aortic rupture and severe clinical consequences^2^. The pathogenesis of AAA is complex, involving multifactorial interactions among genetic, inflammatory and immune factors^3–6^. Despite progress in early diagnosis and treatment, there is still limited understanding of the disease mechanisms, and effective therapeutic strategies remain lacking. Therefore, elucidating the molecular mechanisms underlying AAA, particularly identifying potential therapeutic targets, has become a significant focus in cardiovascular research.

In the formation of AAA, immune-mediated inflammatory responses, metabolic dysregulation^7^, and vascular smooth muscle cell (VSMC) phenotype modulation^8^ play critical roles. Increasing research has identified that post-translational modifications (PTMs), such as acetylation, are pivotal in these biological processes. As a common PTM, acetylation has been shown to play essential roles in regulating cellular function, signal transduction, metabolic pathways, and immune responses^9–12^. In particular, some of the pathological processes in which it is involved also play a key role in AAA.

Recent studies on the role of acetylation in AAA have deepened and revealed potential regulatory networks. For example, acetylation not only regulates the recruitment and activation of inflammatory cells in the vascular wall^13^ but also indirectly affects the mechanical stability of the aortic wall by reshaping VSMC phenotypes and influencing VSMC differentiation, proliferation, and apoptosis^14, 15^. Moreover, acetylation is also important in metabolic regulation, as it modulates the activity of key metabolic enzymes, and impacts the local inflammatory microenvironment^16–18^. These findings suggest that acetylation may be a central regulatory factor in the pathological mechanisms of AAA, providing new theoretical support for further exploration of its pathophysiological mechanisms.

More importantly, histone acetylation is an important epigenetic modification. Studies show that histone acetylation and deacetylation are dynamic and reversible processes, primarily regulated by three protein families: histone acetyltransferases (HATs), histone deacetylases (HDACs), and bromodomain-containing proteins (BRDs). As a key component of epigenetic modifications, histone acetylation plays a critical role in the progression, treatment, and prognosis of many diseases^19–21^. However, systematic and in-depth studies on the specific roles of histone acetylation in different pathological stages and microenvironments of AAA are still lacking. Further analysis is required to unravel the completeness and interaction mechanisms of its molecular network.

In this study, proteomic and acetylomic analyses were performed on abdominal aortic tissues from control and AAA mouse models, leading to the identification of numerous differentially expressed proteins and acetylation sites. Our research findings indicated that histones played an important role in the inflammatory response and immune regulation pathways involved in AAA formation. Notably, a substantial number of acetylated histones involved in this process were regulated by HATs and HDACs. This study aimed to clarify the key regulatory networks of acetylation in AAA development and identified potential therapeutic targets.

## 2 Methods

A detailed description of the methods is provided in the Supplemental Materials.

### 2.1 Animal experiments

All animal treatments and experimental protocols were approved by the Animal Care and Use Committee at the Institute of Basic Medical Sciences, Chinese Academy of Medical Science and Peking Union Medical College (ACUC-A01-2025-006). All animal procedures performed conform to the guidelines from Directive 2010/63/EU of the European Parliament on the protection of animals used for scientific purposes. Mice were maintained on a 12:12 h day: night cycle and provided constant access to food and water.

### 2.2 Mouse model of AAA

Eight-week-old male Apoe knockout (Apoe-/-) mice on a normal chow diet were used in this study. Mice were anesthetized by intraperitoneal injection of avertin at a dose of 187.5 mg/kg body weight, and the surgery was performed only when the toe reflex disappeared. This anesthetic is suitable for experimental scenarios lacking inhalation anesthesia equipment. Its intraperitoneal injection characteristics and rapid onset can meet the surgical requirements under conditions without an inhalation anesthetic machine. This method was conducted in accordance with previously published studies and complied with standard operating procedures^22, 23^. The mice were infused subcutaneously with Angiotensin II (Ang II, Solarbio, A9290) at 1.44 mg/kg/day using osmotic pumps (Alzet, model 2004) for 28 days. Mice in the control group were treated with saline via the same method. The procedure was performed as described in previous studies^22, 23^. In brief, Ang II was dissolved in sterile saline and delivered using Alzet osmotic pumps. Mice were anesthetized with an intraperitoneal injection of avertin, and the pumps were implanted subcutaneously through a small incision at the back of the neck, which was then sutured. The incision healed quickly without infection in any of the mice.

### 2.3 Abdominal ultrasonography in mice

Aortic diameters were measured by high-frequency ultrasound Vevo 2100 echography device (Visual Sonics). Briefly, mice were anesthetized with isoflurane (3%) in oxygen for 2 min until immobile. Each mouse was then taped on a heated (35-37°C) procedure board with isoflurane (1.5%) administered via nosecone on a supine position during ultrasound. Inhalation anesthesia was chosen for the ultrasound examination to maintain hemodynamic stability, which is directly related to the special requirements for the accuracy of cardiac function data in this detection method. The animal ultrasound examination was conducted in a specialized animal ultrasound institution with an inhalation anesthetic machine, so isoflurane was used as the anesthetic. The abdominal area was shaved and coated with ultrasound transmission gel before positioning the acquisition probe. These measurements were performed on the day 0 and day 28 after Ang II infusion. As previously described^23^, long-axis ultrasound scans of suprarenal aortas were performed from the aortic hiatus to the renal artery. Images of the abdominal aortas were obtained. Maximal internal diameters of aortic images were measured with the Vevo LAB software.

### 2.4 Sample collection

At day 28 after Ang II infusion, mice were deep anesthetized with avertin (187.5mg/kg body weight) by intraperitoneal injection. It was used in the animal euthanasia process in this study, which is in line with operational norms. After the toe reflex disappeared, blood was collected and the right atrium was cut open, and saline was perfused through the left ventricle to remove blood in aortas. Then the abdominal aortas were dissected, weighed, and photographed for documentation. After rapid freezing with liquid nitrogen, transfer to -80℃ until proteome and acetylation analysis and detection are performed. Place tissues intended for morphological experiments in 4% paraformaldehyde for fixation for 24 hours, then proceed with paraffin embedding, and store the wax blocks at 4℃ or room temperature until testing.

### 2.5 Proteomic and acetylomic experiments

Samples were retrieved from -80°C and an appropriate amount was weighed into a pre-cooled mortar with liquid nitrogen. Liquid nitrogen was added, and the samples were thoroughly ground into a fine powder. The samples were then collected, lysed with lysis buffer, and digested with trypsin. To enrich the acetylated peptides, the peptides were dissolved in pre-washed resin (antibody resin, catalog number PTM0104, provided by PTM Bio, Hangzhou Jingjie Biotechnology Co., Ltd.). The peptides were separated by a UHPLC system and then injected into an NSI ion source for ionization before being analyzed by Orbitrap Astra mass spectrometry, and the MS/MS data were analyzed using the Spectronaut (v.18) software.

### 2.6 Quantitative Analysis

The raw LC-MS datasets were first searched against database and converted into matrices containing Normalized intensity (the raw intensity after correcting the sample/batch effect) of proteins. The Normalized intensity (I) was transformed to the relative quantitative value (R) after centralization. The formula is listed as follow where i represents sample and j represents protein:

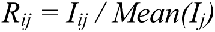

### 2.7 Repeatability Analysis

For experiment with biological or technical replicates, it is necessary to evaluate the quantitative reproducibility among the biological or technical replicates. Three statistical analysis methods including Principal Component Analysis (PCA), Relative Standard Deviation (RSD) and Pearson’s Correlation Coefficient were used to evaluate the quantitative reproducibility.

## 3 Results

### 3.1 Formation of the abdominal aortic aneurysm model in mouse

Six mice were divided into two groups and three mice in each group were respectively treated with saline and angiotensin II (Ang II) for 28 days. Ang Ⅱ stimulation induced abdominal aortic aneurysms in mice **(Fig. 1A)**. Subsequently, it could be observed that compared with the control group, the disease group mice showed a significant increase in systolic blood pressure **(Fig. 1B)** and diastolic blood pressure **(Fig. 1C)**, while the heart rate **(Fig.1D)** was basically the same. Meanwhile, aorta weight/body weight **(Fig. 1E)** was significantly increased and the ultrasound images revealed an enlargement in the diameter of the abdominal aorta **(Fig. 1F, and 1G)**. The above results indicated that abdominal aortic aneurysms had been successfully formed in mice.

**Figure 1.**
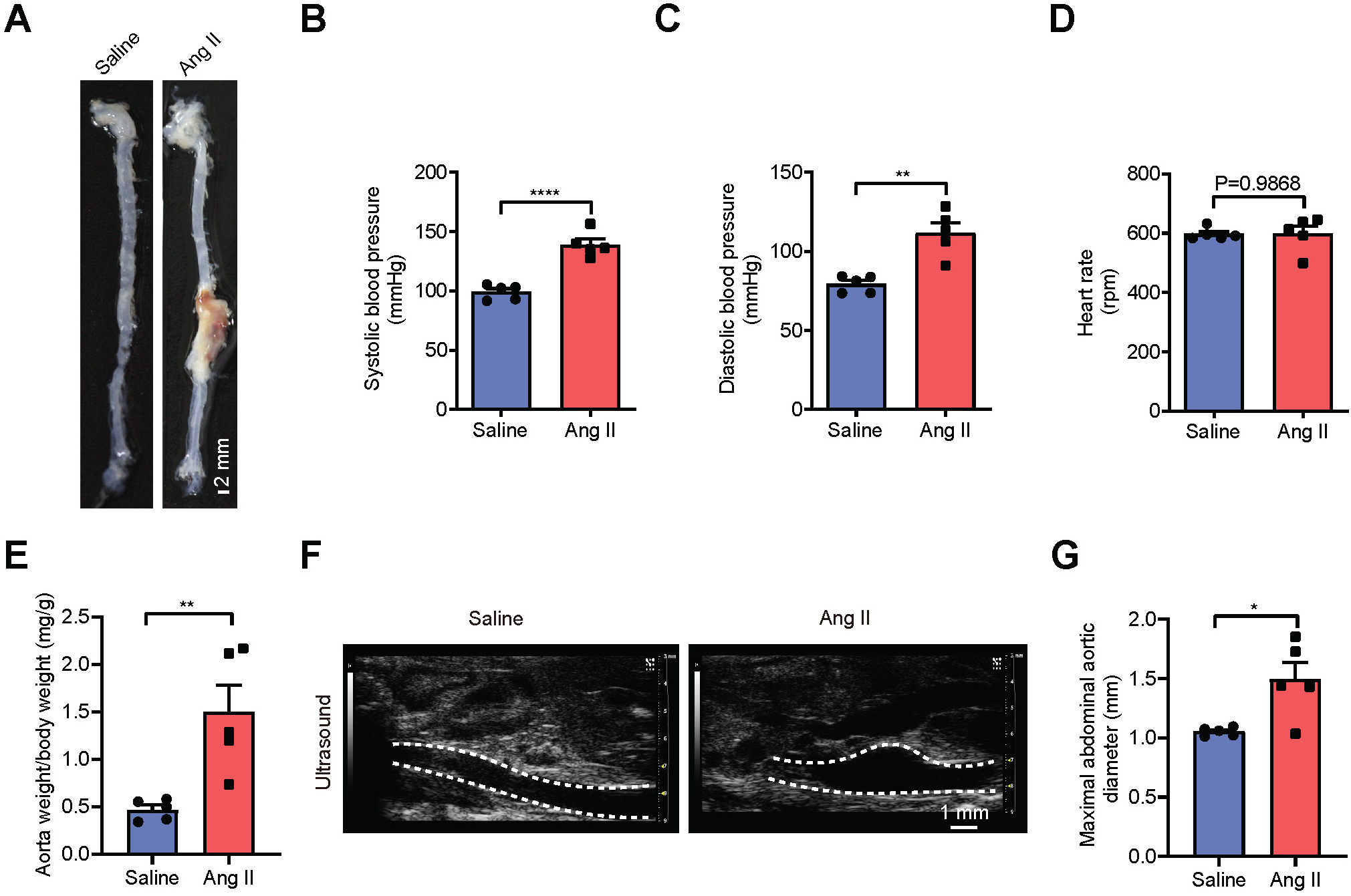
Formation of the abdominal aortic aneurysm model in mice. **(A).** Representative images (scale: 2 mm) showing macroscopic features of the normal aorta and aneurysms in mice after infusion with saline (n=3) or Ang II for 28 days (n=3). **(B-E).** Changes in physiological indicators included systolic blood pressure (B), diastolic blood pressure (C), heart rate (D) and aorta weight/body weight (E) between the control and AAA groups after 28 days of infusion with saline and Ang II. **(F).** Ultrasound images of abdominal aortic in mouse (scale: 1 mm). **(G).** Differences in the maximum diameter of the abdominal aorta.

### 3.2 Global profiling of the proteome and acetylome

We performed label-free proteome and acetylome analysis on aortic tissue samples from six mouse (three from UT and three from AngⅡ_28d group). The workflow for the study was presented in **Fig. 2A**, and the detailed methods are listed in the Supporting Information. Mass spectrometry (MS) raw files were analyzed using Spectronaut (v.18) software. A total of 7812 and 7852 quantifiable proteins were separately identified in the proteomes **(Fig. 2B and S1A)**. This number exceeded the numbers of proteins previously detected in AAA tissues of mice^24^. In the acetylome, 4497 lysine acetylation (Kac) sites were identified in the control group and 4565 in the AAA group **(Fig. 2C and S1B)**. The Pearson correlation coefficients in **Fig. 2D and 2E** indicated high consistency between biological replications. Principle component analysis revealed that proteins **(Fig. 2F)** and Kac sites **(Fig. 2G)** of the two groups were located in different clusters, clearly distinguishing the UT and AngⅡ_28d models.

**Figure 2.**
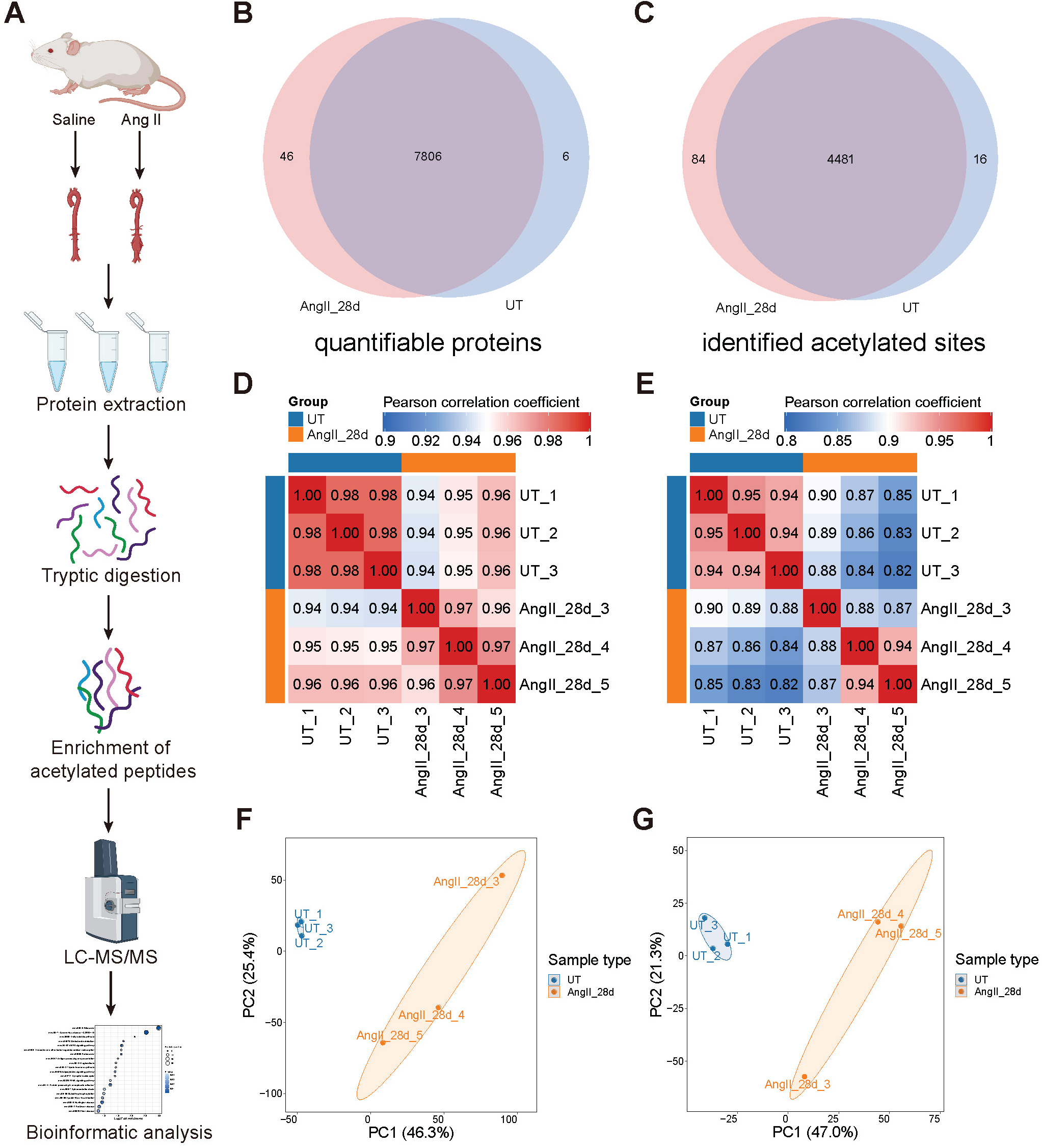
Comprehensive analysis workflow of protein and acetylation Features. **(A).** The flow chart depicted the experimental procedures and data analysis workflow (Created in https://BioRender.com). **(B, C).** Venn diagrams showed the overlapped features including 7806 quantifiable proteins (B) and 4481 Kac sites (C) among proteome and acetylome. **(D, E).** Pearson correlation analysis of quantifiable proteins (D) or Kac sites (E) between two groups of samples (AngⅡ_28d and UT, n = 3 per group). **(F, G).** Principal component analyses visualizing the sample distribution according to their proteome (F) and acetylome profiles (G).

### 3.3 Differential proteins and functional enrichment analyses of proteome

Based on proteomic analysis of the UT and AngⅡ_28d group, 7858 proteins were quantified, and 1048 proteins were defined as significantly regulated proteins **(Table S1**). Among them, 576 proteins were significantly upregulated, while 472 were significantly downregulated **(Fig. 3A and S2A)**. The heatmaps showed differences in the expression of significantly regulated proteins between UT and AngⅡ_28d group **(Fig. 3B)**. To understand the functions of differential proteins and their impact on biological processes, a Kyoto Encyclopedia of Genes and Genomes (KEGG) pathway enrichment analysis was conducted for differential proteins **(Table S2, S3 and S4)**. We found that upregulated pathways were related to immunological disorders and downregulated pathways were related to cardiological diseases **(Fig. 3C, S2B and S2C)**, such as complement and coagulation cascades, systemic lupus erythematosus (SLE), arrhythmogenic right ventricular cardiomyopathy (ARVC), dilated cardiomyopathy (DCM), hypertrophic cardiomyopathy (HCM). Chord diagram **(Fig. 3D)** depicted the containment relationship between up and down regulated each Top 5 pathways and proteins.

**Figure 3.**
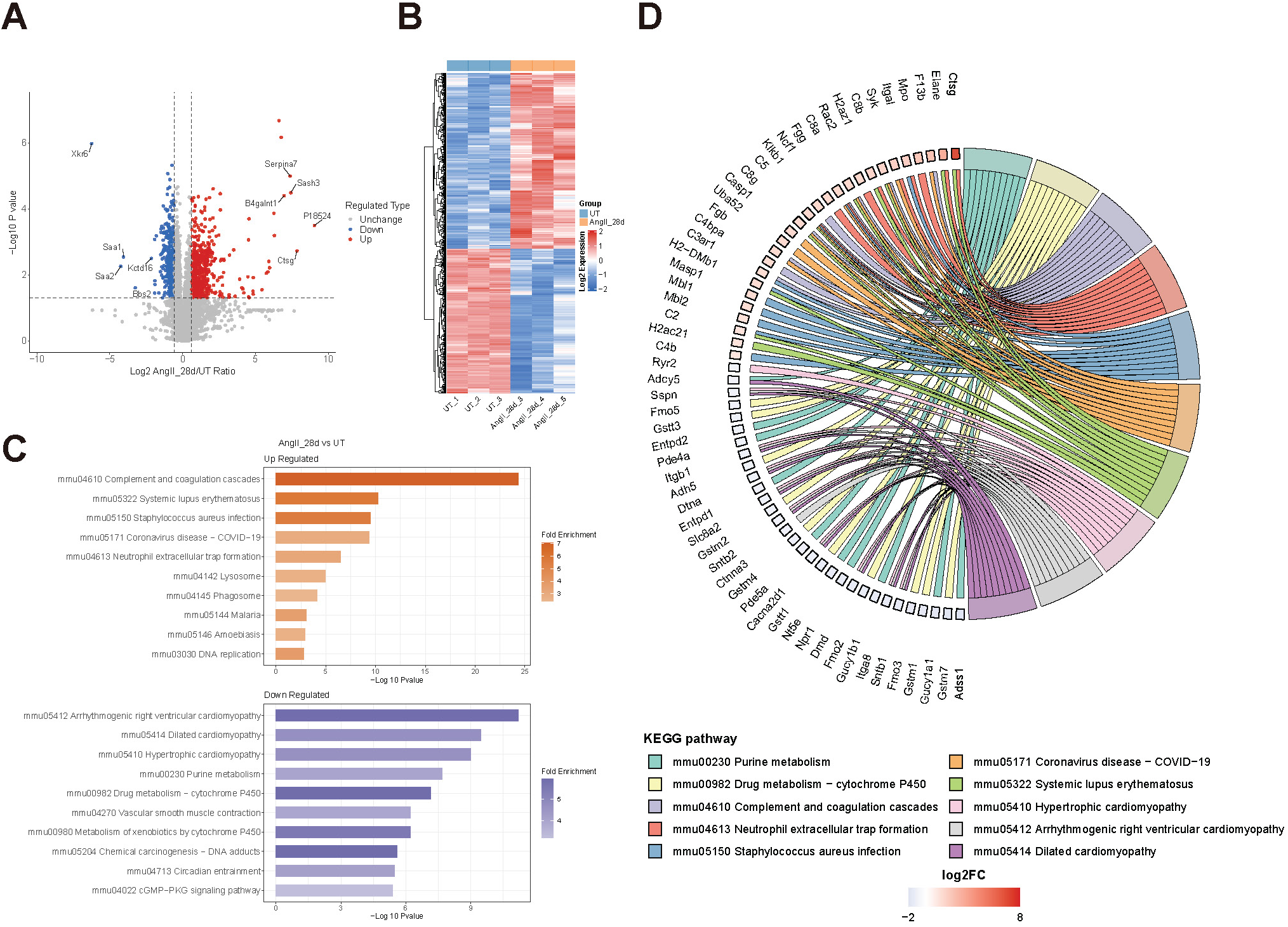
Differential proteins and functional enrichment analyses of proteome. **(A).** Volcano plot displayed the differential proteins between AngⅡ_28d (n=3) and UT (n=3) groups. Red dots represented 576 significantly upregulated quantifiable proteins (T test, p value < 0.05, fold change > 1.5) and blue dots represented the 472 downregulated quantifiable proteins (T test, p value < 0.05, fold change < 0.6667). **(B).** The heatmap showing the relative expression levels of differentially expressed proteins in UT and AngⅡ_28d groups. **(C).** Top 10 enriched items identified from the Kyoto Encyclopedia of Genes and Genomes (KEGG) pathway enrichment analysis of up and down regulated proteins (Fisher’s exact test, P value <0.5). **(D).** Chord diagram depicted the inclusion relationship of pathways and proteins.

### 3.4 Differential proteins and functional enrichment analyses of acetylome

We detected and defined 1184 significantly regulated acetylation sites **(Table S5)**. Among them, 503 sites were significantly upregulated, while 681 were significantly downregulated **(Fig. 4A and S3A)**. The heatmaps showed differences in the expression of significantly regulated proteins between UT and AngⅡ_28d group **(Fig. 4B)**. Subsequently, KEGG enrichment analysis were performed **(Table S6, S7 and S8)**, and some of the pathways upregulated at the differential sites were also associated with inflammatory responses and immunological disorders **(Fig. 4C and S3B)**. Moreover, downregulated pathways were related with metabolic pathways **(Fig. 4C and S3C)**, such as valine, leucine and isoleucine degradation, lysine degradation, and propanoate metabolism. Chord diagram depicted the containment relationship between up and down regulated each Top 5 pathways and sites **(Fig. 4D)**.

**Figure 4.**
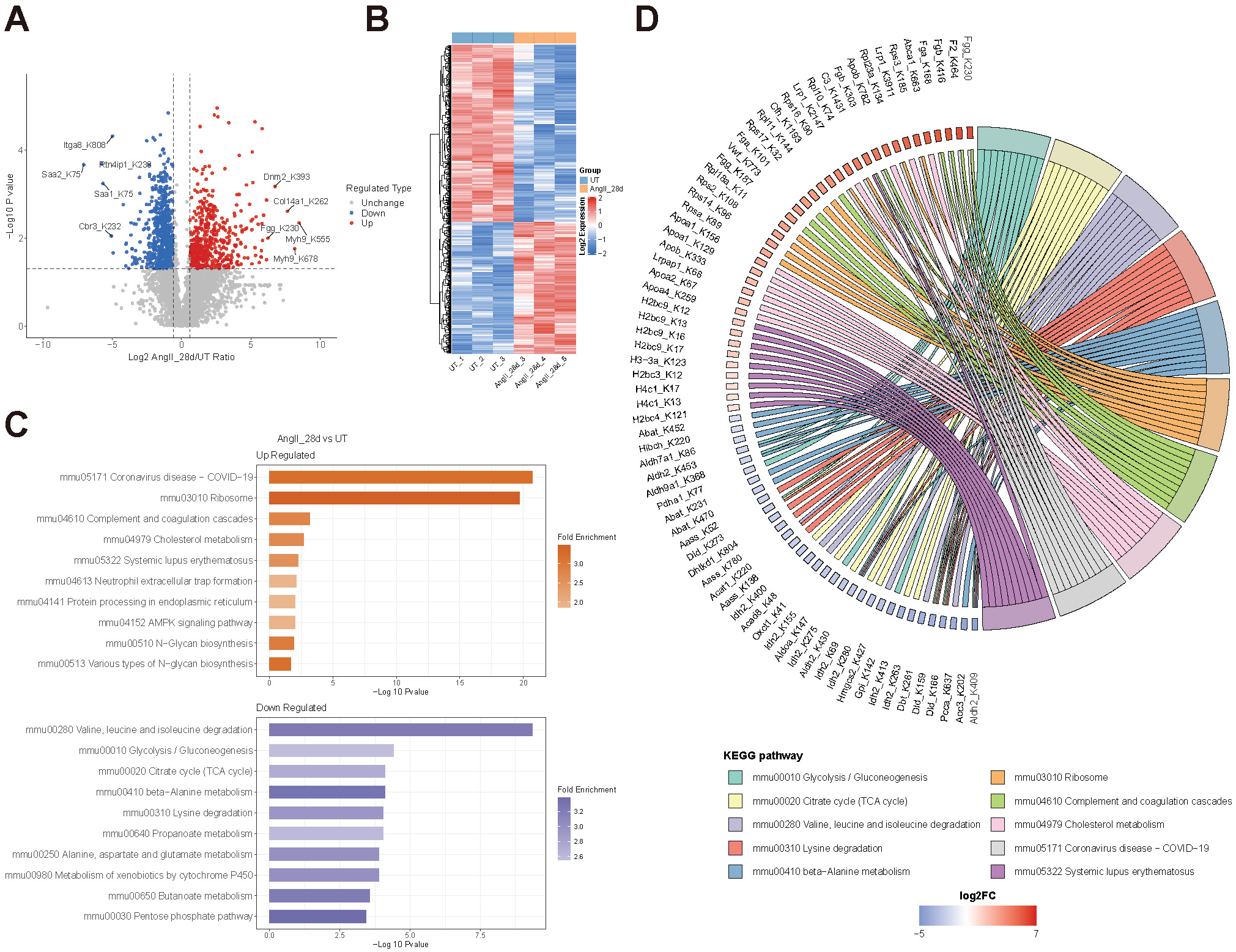
Differential proteins and functional enrichment analyses of acetylome. **(A).** Volcano plot displayed the differential Kac sites between AngⅡ_28d (n=3) and UT (n=3) groups. Red dots represented 503 significantly upregulated quantifiable Kac sites (T test, p value < 0.05, fold change > 1.5) and blue dots represented the 681 downregulated quantifiable Kac sites (T test, p value < 0.05, fold change < 0.6667). **(B).** The heatmap showing the relative expression levels of differentially expressed Kac sites in UT and AngⅡ_28d groups. **(C).** Top 10 enriched items identified from the KEGG pathway enrichment analysis of up and down regulated acetylated sites (Fisher’s exact test, P value <0.5). **(D).** Chord diagram depicted the inclusion relationship of pathways and sites.

### 3.5 Correlation analysis of proteome and acetylome

By correlation analysis of quantified proteins with quantified acetylation-modified proteins, we found that 1721 proteins appeared in both the proteome and the acetylome **(Fig. 5A)**. The correlation analysis between quantifiable proteins and acetylation quantifiable sites revealed a low correlation between protein abundance and acetylation levels **(Fig. 5B)**. This may suggest that acetylation is regulated by other mechanisms, such as the activity of specific HATs or HDACs, rather than solely depending on protein expression levels. We performed “Nine-Square scatterplot” analysis of the quantitative proteome and acetylome data **(Table S9)**. The proteins were divided into nine groups **(Fig. 5C)**. Subsequently, KEGG functional enrichment analysis was performed on the proteins in each quadrant **(Fig. S4A-F)**, and the both upregulated pathways of proteins and acetylated proteins were selected for bubble mapping **(Fig. 5D)**. It is indicated that the same KEGG pathways for proteome and acetylome included Complement and coagulation cascades, Systemic lupus erythematosus (SLE), Coronavirus disease − COVID−19 and NET formation. Protein-protein interaction networks (PPI) among these KEGG pathways were performed via the STRING website (https://string-db.org/) and Cytoscape (v.3.10.3) software **(Fig. 5E)**. These pathways are all related to the inflammatory response, and we found a significant number of acetylated histones, suggesting that histones may play an important role in AAA. Therefore, we further explored the histone acetylation process.

**Figure 5.**
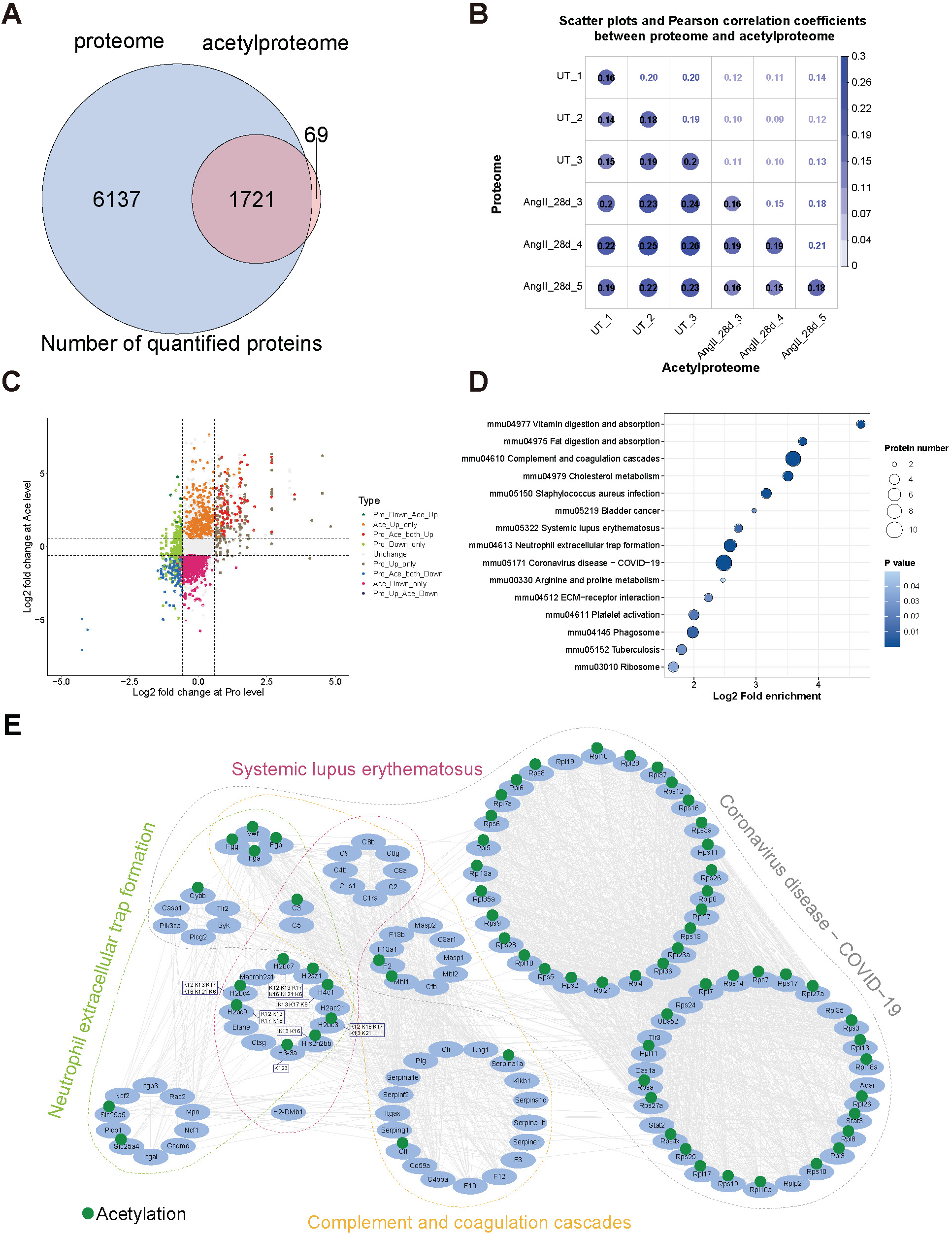
Correlation analysis of proteome and acetylome. **(A).** The Venn diagram illustrated the overlap of proteins identified in the proteome and acetylome. **(B).** Pearson correlation coefficients between proteome and acetylome. **(C).** The nine-quadrant scatterplot for fold changes of proteins and acetylation proteins. **(D).** KEGG enrichment bubble plot of upregulated Proteins in both proteome and acetylome (Fisher’s exact test, P value <0.5). **(E).** Protein-protein interaction (PPI) networks of identified proteins in the proteome and acetylated proteins in the acetylome were involved in the Complement and coagulation cascades, Systemic lupus erythematosus, Coronavirus disease − COVID−19 and Neutrophil extracellular trap formation. The proteins identified in the proteome and acetylome that were highlighted in blue circles and green dots were determined to be modified by acetylation in this study, and the boxes indicated partly the acetylation sites (STRING database 11.5, confidence score > 0.4).

### 3.6 Global analysis of histone acetylation

Histone acetylation is a central element in cellular regulatory networks and is important for understanding the regulation of gene expression, cell fate decisions, and disease mechanisms^25^. We identified numerous histones and acetylated histones **(Table S10 and S11)**, then counted the number of Kac sites occurring on the lysine of H1 linker proteins and core histones H2A, H2B, H3, and H4 **(Fig. 6A)**, with the highest number of modifications occurring in H2B. Subsequently, we compared the acetylation levels of H1, H2A, H2B, H3, and H4 as a whole between the two groups, and the results showed that none of them differed significantly **(Fig. 6B)**. Then, we compared the acetylation levels of specific histone modification sites in both AngII_28d/UT groups **(Fig. 6C)**, of which 28 sites were significantly different and all were upregulated in the AngII_28d group, and most of these sites occurred in the H2B histone. As acetylation is reversible and related to acetyltransferases, deacetylases, therefore we detected the protein intensity of differentially acetylation-related enzymes, BRDs and histone in both AngII_28d/UT groups (Figure 6D and Table S12, S13, S14). The results indicated that BRD, acetyltransferases, and deacetylases were all downregulated. Meanwhile, histones were all upregulated. We did a Pearson correlation of the differential histone modification sites in **Fig. 6C** with the proteins in **Fig. 6D** and plotted the correlation heatmap **(Fig. 6E)**. We found positive associations between Amdhd2 protein levels and several H2B acetylation sites, as well as negative associations between Sirt5, Sirt2 protein levels and most histone acetylation sites levels. This suggested that these deacetylases could regulate histone sites acetylation levels. To demonstrate which histone sites were most affected by the key modifying enzymes, selected histone modification sites and acetylation-related enzymes with significant correlations, and plotted the scatterplots **(Fig. 6F)**. The sites significantly correlated with both sirt2 and sirt5 included Hist2h2bb-K13/K16, H2bc3-K12/K16/K17/K13/K21, H2bc4-K12/K13/K17/K16, H2bc7-K12/K13/K17/K16, H2bc9-K12/K13/K16/K17, H3-3a-K123, and H4c1-K13/K17. Additionally, sirt5 was also correlated with H1-1-K121, H2bc4-K121/K6, H2bc7-K121/K6, and H4c1-K9. It can be seen that the majority of histone sites are significantly regulated by deacetylases, which influence key pathways in AAA by modulating histone acetylation. These may serve as new therapeutic targets for AAA.

**Figure 6.**
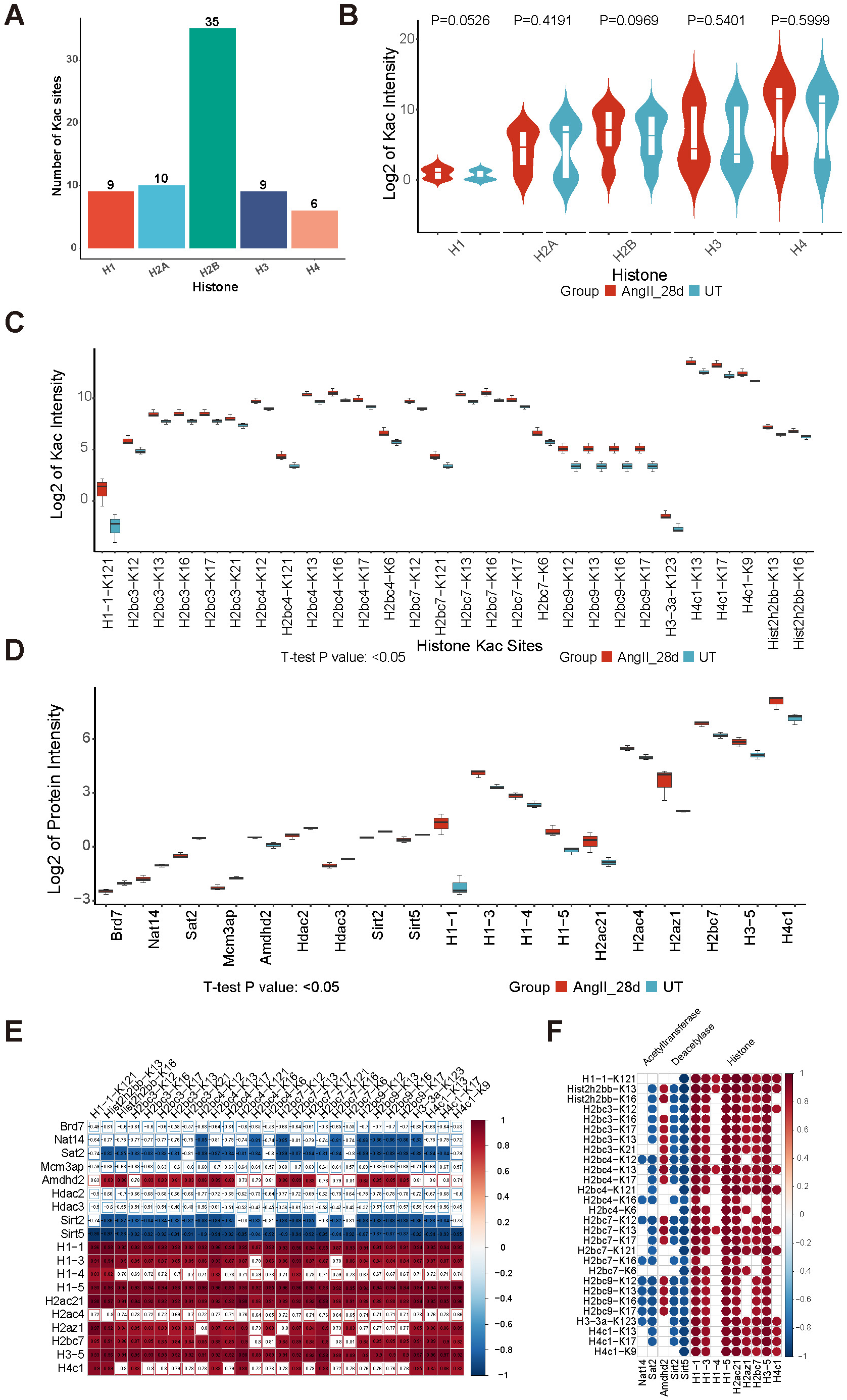
Global analysis of histone acetylation. **(A).** Number of Kac sites on histone H1 H2A, H2B, H3, H4. **(B).** Violin plot compared acetylation levels of H1, H2A, H2B, H3, H4 as a whole in the AngII_28d and UT groups (n = 3 per group, and T test p value). **(C).** Box plots demonstrated the level of acetylation of specific histone modification sites in both AngII_28d and UT groups (T test, p value < 0.05). **(D).** Box plots showed protein intensity of acetylation-associated enzymes, BRDs, histone in both AngII_28d/UT groups (T test, p value < 0.05, fold change > 1). **(E).** Heatmap of Pearson correlation between differential histone modification sites in C and proteins in D (unfilled colors with P value > 0.05). **(F).** Scatterplot demonstrated interactions between histone modification sites and acetylation-associated enzymes with significant correlations in E.

## 4 Discussion

In this study, we performed a differential analysis of proteins and acetylation modifications in the abdominal aortic tissues of control and AAA mouse models, and identified a large number of proteins and acetylation sites. KEGG enrichment analysis revealed many pathways related to inflammation and immunity response both in the proteome and acetylome (Figure 3C, 4C). AAA is a chronic inflammatory disease primarily associated with extracellular matrix (ECM) destructive remodeling, VSMC apoptosis, inflammation, and oxidative stress^26–28^. These pathways also suggest that immune cells and inflammatory factors play an important role in the progression of AAA. Immune cells contribute to aortic dilation and rupture by secreting inflammatory factors IL-1β and IL-18 that promote apoptosis of aortic wall cells and drive the phenotypic transition of VSMC^29, 30^. Additionally, they release proteases that degrade the ECM, further weakening the aortic structure. Neutrophils, the most abundant and rapidly responding immune cells in the human immune system, can exacerbate AAA through the activation of matrix metalloproteinases (MMPs) and the inactivation of their inhibitors TIMP^31^. The involvement of NET is crucial in the formation of AAA^32^, as NETs activate NLRP3 in macrophages, further releasing IL-1β and IL-18, which upregulate inflammation in the aorta.

Additionally, we found differential expression in acetylation-regulated metabolic pathways, with cholesterol metabolism pathways upregulated, while amino acid metabolism, glycolysis, and TCA cycle pathways were downregulated (Figure 4C). Cholesterol and its derivatives (such as oxidized cholesterol) are important molecules in inflammation regulation^33^. Inflammatory factors like TNF-α, IL-6, and IL-1β can also can also affect lipid metabolism. Upregulated cholesterol metabolism supports the generation of lipid signaling molecules, amplifying lipid peroxidation and inflammatory signaling, such as activating macrophages via oxidized cholesterol, further promoting inflammation and immune cell function^34^. Oxidized cholesterol also accelerates the degradation of the vascular wall by participating in MMP activation^35^, further advancing the pathological progression of AAA. The inhibition of amino acid metabolism, glycolysis, and TCA cycle reflects the shift in metabolic priorities from energy production to inflammatory signaling in the inflammatory environment^36, 37^. The mitochondria in the vascular wall of AAA patients are usually damaged by oxidative stress and inflammation, leading to decreased TCA cycle efficiency and accumulation of glycolytic intermediates.

We performed PPI analysis on the pathways that were upregulated in both the proteome and acetylome, including Complement and coagulation cascades, SLE, Coronavirus disease − COVID−19, and NET formation. The complement system is an important component of the innate immune system, involved in processes such as inflammation, vascular remodeling, and thrombosis. In the onset and development of AAA, the complement system promotes local inflammatory responses and the degradation of the vessel wall by activating the complement cascade, thereby influencing the formation and progression of AAA^38^. Systemic lupus erythematosus is an autoimmune disease characterized by abnormal activation of the immune system, leading to inflammation and damage to the blood vessel walls. The development of AAA is also associated with chronic inflammation of the vessel wall, where the activation of immune cells and the secretion of cytokines play important roles in the pathological processes of both conditions. Coronavirus binds to the ACE2 receptor on host cells through its spike glycoprotein, potentially causing endothelial cell damage and dysfunction, thereby increasing the risk of vascular diseases. The NET formation pathway is closely related to AAA. Studies have shown that blocking thrombosis-dependent NET can reduce the progression of AAA^32, 39^. Inhibiting the formation or function of NETs may serve as a new strategy for the treatment of AAA. we noticed that there were abundant acetylated histones in the NETs formation pathway. Histone acetylation, through regulating chromatin structure and gene expression, plays a widespread and crucial role in cellular functions and life processes^19–21^. This dynamic modification mechanism is not only a switch for gene expression but also involves processes like metabolism, development, immunity, and DNA repair. The acetylation modification sites on histone H2B were the most abundant, likely due to the highly exposed lysine residues. The N-terminal tail of histone H2B has multiple lysine residues that are exposed on the surface of chromatin, making them major targets for acetylation. A total of 69 differential histone modification sites were identified (Figure 6A), with 28 showing significant differences, all upregulated in the AAA group (Figure 6C). Since acetylation is reversible and associated with acetyltransferases and deacetylases, we examined the abundance of acetylation-related enzymes, BRDs, and histones in the AngII_28d/UT groups (Figure 6D). We found that histone deacetylases were downregulated, and the sirtuin family proteins sirt2 and sirt5 were significantly negatively correlated with most differential acetylation sites (Figure 6E). The sirtuin family of proteins has an important role in immune regulation and inflammatory responses^40–43^, and a number of studies have reported its effects in inhibiting aortic aneurysms ^22, 23, 44–46^. We speculate that promoting the expression of Sirt2 and Sirt5 may reduce histone acetylation levels, potentially inhibiting the formation or function of NETs. This could in turn suppress immune and inflammatory processes in AAA, thereby limiting its progression and making it a potential therapeutic target for AAA.

## Conclusions

In summary, this study provides comprehensive proteomic and acetylomic data for the abdominal aortic tissue in the infected mouse model of aortic aneurysm, significantly expanding the existing acetylation database in mice. It emphasizes the importance of histones in inflammation response and immune dysregulation pathways. The results suggest that sirt2 and sirt5 may potentially modulate immune and inflammatory processes in the NETs, becoming potential targets for AAA progression and treatment, and also offer new possibilities for exploring therapeutic targets within the sirtuin protein family. However, our study also has some limitations, such as the differences between mouse and human aortic tissues and the lack of experimental validation. Further research is needed to confirm the effectiveness of these targets.

## Data availability statement

The raw proteome and acetylome mass spectrometric data had been deposited to the ProteomeXchange Consortium via the iProX partner repository (https://www.iprox.org/) with the dataset identifiers PXD060895^47, 48^.

URL: https://proteomecentral.proteomexchange.org/cgi/GetDataset?ID=PXD060895

## Acknowledgements

We thank the Chinese Academy of Medical Sciences & Peking Union Medical College and all authors who contributed to the article.

## Supplemental Material

Supplemental Methods

Figure S1-S4

Tables S1–S14

Major Resources Table

## Author contributions

Luyao Zhang analyzed the data and wrote the manuscript; Bo Yang conducted the experiments and analyzed the data; Tao Ding, Yaling Zhao, Yehong Yang, Xiaoyue Tang, Yue Wu, Qiaochu Wang, Zhiyi Zhang, Chunmei Shi, Ron Han, and Xutong Zhang analyzed the data and provided valuable discussions; Jiangfeng Liu, Juntao Yang performed data management and directed and designed the experiments. All authors gave final approval of the submitted version and agreed to take responsibility for all aspects of the work.

## Highlights

- Comprehensive proteomic and acetylomic analyses identified 7,858 proteins and 1,790 acetylated proteins in AAA mouse aortic tissue.
- Bioinformatics revealed that histone modifications are central to inflammation and immune dysregulation in AAA.
- Sirt2 and Sirt5 were identified as potential regulators of neutrophil extracellular trap (NET) formation via histone deacetylation.
- Findings highlight the therapeutic potential of targeting the sirtuin family for AAA treatment.

